# It’s not you, it’s me: Corollary discharge in precerebellar nuclei of sleeping infant rats

**DOI:** 10.1101/329540

**Authors:** Didhiti Mukherjee, Greta Sokoloff, Mark S. Blumberg

**Affiliations:** Department of Psychological and Brain Sciences, University of Iowa, Iowa City, Iowa; Interdisciplinary Graduate Program in Neuroscience, University of Iowa, Iowa City, Iowa; Department of Biology, University of Iowa, Iowa City, Iowa; Iowa Neuroscience Institute, University of Iowa, Iowa City, Iowa; Delta Center, University of Iowa, Iowa City, Iowa

## Abstract

In week-old rats, somatosensory input arises predominantly from stimuli in the external environment or from sensory feedback associated with myoclonic twitches during active (REM) sleep. A previous study of neural activity in cerebellar cortex raised the possibility that the brainstem motor structures that produce twitches also send copies of motor commands (or corollary discharge, CD) to the cerebellum. Here, by recording from two precerebellar nuclei—the inferior olive and lateral reticular nucleus—we demonstrate that CD does indeed accompany the production of twitches. Within both structures, the CD signal comprises a surprisingly sharp activity peak within 10 ms of twitch onset. In the inferior olive, this sharp peak is attributable to the opening of slow potassium channels. We conclude that a diversity of neural activity is conveyed to the developing cerebellum preferentially during sleep-related twitching, enabling cerebellar processing of convergent input from CD and reafferent signals.

## Introduction

The sensorimotor systems of diverse vertebrate and invertebrate species distinguish signals arising from self-generated movements (i.e., reafference) from those arising from other-generated movements (i.e., exafference; Cullen, 2004). To make this distinction, motor structures generate copies of motor commands, referred to as corollary discharge (CD; Crapse and Sommer, 2008; Poulet and Hedwig, 2007). CD is conveyed to non-motor structures to inform them of the imminent arrival of reafference arising from self-generated movements (Crapse and Sommer, 2008). By comparing the two signals, animals are able to distinguish between self-produced and other-produced movements.

Self-produced movements are not restricted to periods of wakefulness, especially during development. Infants produce brief, discrete, jerky movements of skeletal muscles during active sleep (AS or REM sleep), which is the predominant behavioral state during early infancy (Jouvet-Mounier et al., 1970; Roffwarg et al., 1966). These spontaneous movements are known as myoclonic twitches, and are among the most conspicuous behaviors during development in a diversity of species (Blumberg et al., 2013; Gramsbergen et al., 1970; Jouvet-Mounier et al., 1970; Roffwarg et al., 1966).

It was recently reported in week-old rats that the external cuneate nucleus (ECN), a medullary nucleus that receives proprioceptive input from the forelimbs, actively inhibits reafference arising from wake-related limb movements but not those arising from twitches (Tiriac and Blumberg, 2016). This state-dependent gating by the ECN suggested the selective engagement of a CD mechanism during wake movements and its suspension during twitching. It remained unclear, however, whether the suspension of the gating mechanism reflected the absence of a twitch-related CD signal or the inhibition of CD’s effects within the ECN. Determining whether twitches are accompanied by CD has implications for their functional role in the self-organization of the sensorimotor system.

There is some suggestive evidence that twitches are accompanied by CD. In week-old rats, twitches trigger both complex spikes (arising from climbing fibers) and simple spikes (arising from mossy fibers) in cerebellar Purkinje cells (Sokoloff et al., 2015a). These neural events were detected at very short latencies that appear too short for reafferent signals arising from the periphery (Puro and Woodward, 1977a; Puro and Woodward, 1977b). Accordingly, it is possible that the motor structures that produce twitches also convey CD to the cerebellum, as occurs with waking movements in adults (Azim and Alstermark, 2015; Azim et al., 2014).

If twitch-related CD reaches cerebellar cortex, it must be conveyed through the precerebellar nuclei. The inferior olive (IO) is a good candidate structure for such a CD signal. First, it is the sole source of climbing fibers to cerebellar cortex and is therefore responsible for the triggering of complex spikes (Ruigrok et al., 2014). Second, midbrain motor structures, including Nucleus of Darkschewitsch, project directly to the IO (De Zeeuw et al., 1998). Finally, the IO fires precisely at the onset of self-generated movements in waking adults (Keating and Thach, 1995; Welsh et al., 1995).

With respect to mossy fibers, there are a few major candidate structures to consider (Ruigrok et al., 2014). First, the pontine nucleus is an unlikely source of CD in week-old rats because it receives descending input from motor cortex (Lee and Mihailoff, 1990), which does not contribute to the production of twitches (Blumberg, 2010; Kreider and Blumberg, 2000). Second, the ECN can also be ruled out as a source of CD because it has been demonstrated to process twitch-related reafference exclusively (Tiriac and Blumberg, 2016). Finally, the LRN is a viable candidate because it receives both sensory input from the limbs and motor input from midbrain structures, including the red nucleus (Alstermark and Ekerot, 2013; Pivetta et al., 2014). Moreover, the LRN has been implicated in processing CD associated with self-generated movements in adults (Alstermark and Ekerot, 2015; Arshavsky et al., 1978).

Accordingly, we recorded neural activity in the IO and LRN in postnatal day (P) 7-9 (hereafter P8) rats. In the majority of IO units and a subset of LRN units, twitch-related activity was remarkably precise, exhibiting a sharp activity peak within ±10 ms of a twitch. This sharp peak, centered so tightly around twitch onset, is most parsimoniously explained as CD. Next, we identified adjacent, non-overlapping midbrain motor areas that are involved in the production of twitches and that are likely sources of CD to the IO and LRN. Finally, we demonstrate that calcium-activated slow potassium (SK) channels in the IO are responsible for sharpening the broad CD signal arriving from motor structures.

Altogether, the twitch-related activity peaks in the IO and LRN reported here satisfy the major criteria put forward for identifying CD (Poulet and Hedwig, 2007; Sommer and Wurtz, 2008). To our knowledge, these findings provide the first direct neurophysiological evidence of a CD signal in an infant mammal.

## Results

### IO activity predominates during active sleep

We recorded IO activity in unanesthetized head-fixed pups as they cycled spontaneously between sleep and wake with their limbs dangling freely (n=20 pups, 37 units, 1-4 units/pup). Electromyography (EMG) and behavioral scoring were used to identify behavioral state and detect sleep and wake movements (Blumberg et al., 2015; Figure 1A).

**Figure 1.**
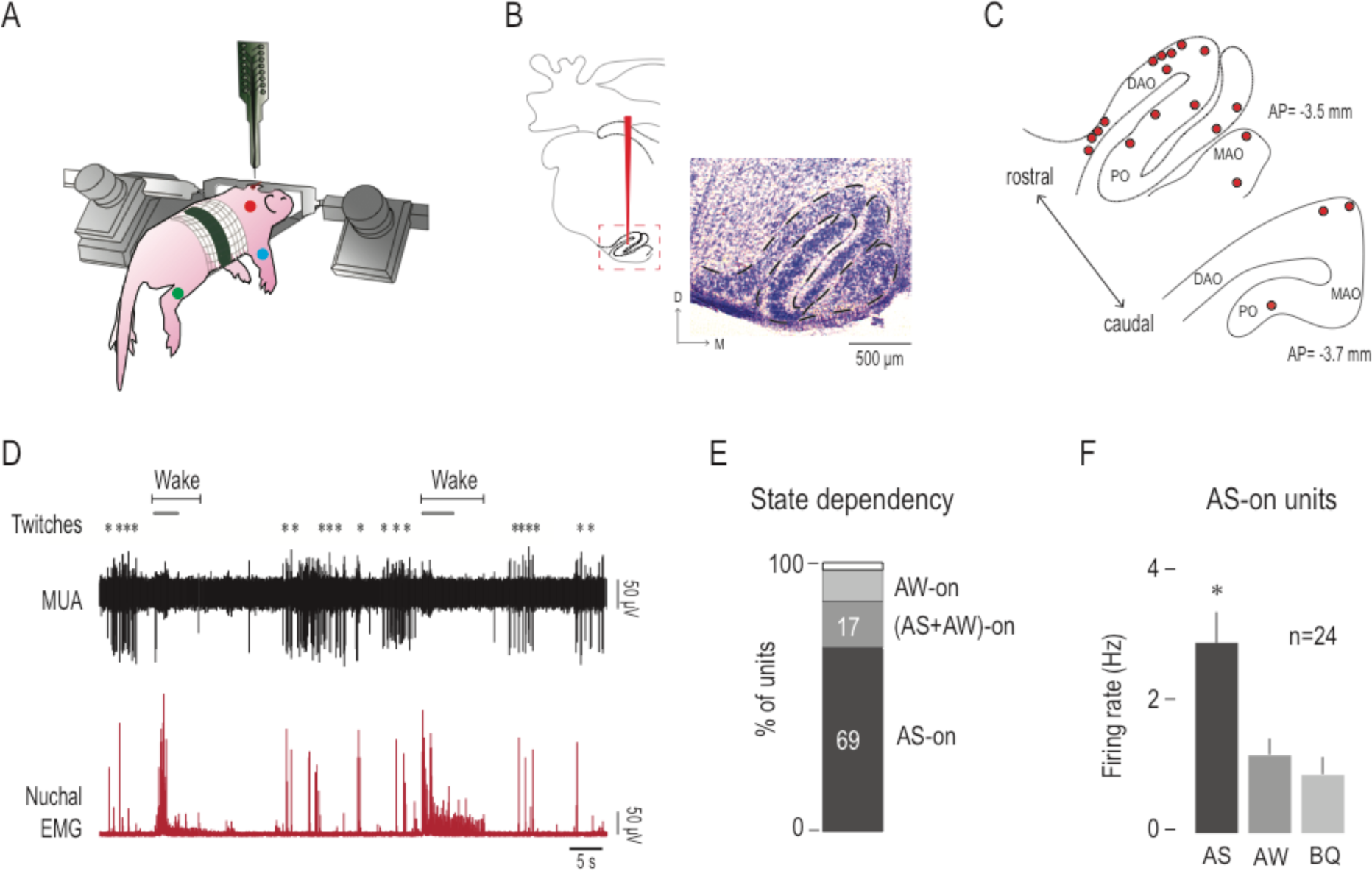
Olivary activity predominates during active sleep. (**A**) Illustration of a head-fixed rat pup in a recording apparatus instrumented with nuchal (red), forelimb (blue), and hindlimb (green) EMG electrodes. (**B**) Left: Reconstruction of a representative electrode placement within the IO (red vertical line). Red dashed line circumscribes the IO. Right: Coronal section stained with cresyl violet depicting the anatomical location of the IO (black dashed line). (**C**) Reconstruction of electrode placements (red circles) within the IO in two coronal sections across all subjects. (**D**) Representative recording of rectified nuchal EMG activity and multiunit activity (MUA) in the IO during spontaneous sleep-wake cycling. Asterisks denote twitches and gray horizontal bars denote wake movements as scored by the experimenter. (**E**) Stacked plot depicting the percentage of units that were AS-on, (AS+AW)-on, and AW-on units in the IO. (**F**) Bar graphs showing mean (+SEM) firing rates of AS-on units across behavioral states. * significant difference from AW and BQ, p<0.0005. D=dorsal; M=medial; DAO=dorsal accessory olive; MAO=medial accessory olive; PO=principal olive; AP=antero-posterior in relation to lambda; AS=active sleep; AW=active wake; BQ=behavioral quiescence.

Electrode placement within the IO was confirmed histologically (Figure 1B). Recording sites were located within the dorsal accessory olive (DAO; n=19 units across 12 pups) or the medial accessory olive (MAO) and the principal olive (PO; n=18 units across 8 pups; Figure 1C). Overall, unit activity was phasic and largely restricted to periods of AS; also, activity appeared to be suppressed immediately after the onset of active wake (AW). Sparse activity was observed during behavioral quiescence (BQ), which is a period of low muscle tone interposed between AW and AS (Figure 1D). The majority (24/35 units, 69%) of IO units were AS-on (Figure 1E) and the mean firing rate of the AS-on units (2.9 ± 0.4 Hz) was approximately three times higher during AS than for the other two states (p<0.0005; Figure 1F). Two IO units were excluded from state analysis due to movement artifact during AW.

### IO neurons exhibit sharp activity peaks at twitch onset

The phasic IO activity clustered around periods of myoclonic twitching; therefore, we examined the temporal relationship between twitches and unit activity by creating perievent histograms (5-ms bins, 1-s window) with unit activity triggered on twitch onset.

Previous studies have revealed two distinct patterns of twitch-triggered perievent histograms in sensorimotor structures (Figure 2A). First, in a motor structure like the RN (green trace in Figure 2A), unit activity increases 20-40 ms before the onset of a twitch (Del Rio-Bermudez et al., 2015). Second, in a sensory structure like the ECN (blue trace in Figure 2A), unit activity increases at least 10-50 ms after the onset of a twitch (Tiriac and Blumberg, 2016). In the IO, however, we found that the majority of units (23/37 units, 62%) were active within +10 ms of the onset of a twitch (Figure 2B, C). These IO activity profiles were very sharp and thus strikingly different from those observed in any of the motor and sensory structures from which we have previously recorded (e.g., see Figure 2D). Also, the IO units that exhibited this profile were responsive primarily to nuchal and/or forelimb twitches and rarely to hindlimb twitches (Figure 2—figure supplement 1A-D). Finally, the characteristics of the neural responses recorded in the IO did not appear to differ across anatomical subdivisions.

**Figure 2 and 1 supplement.**
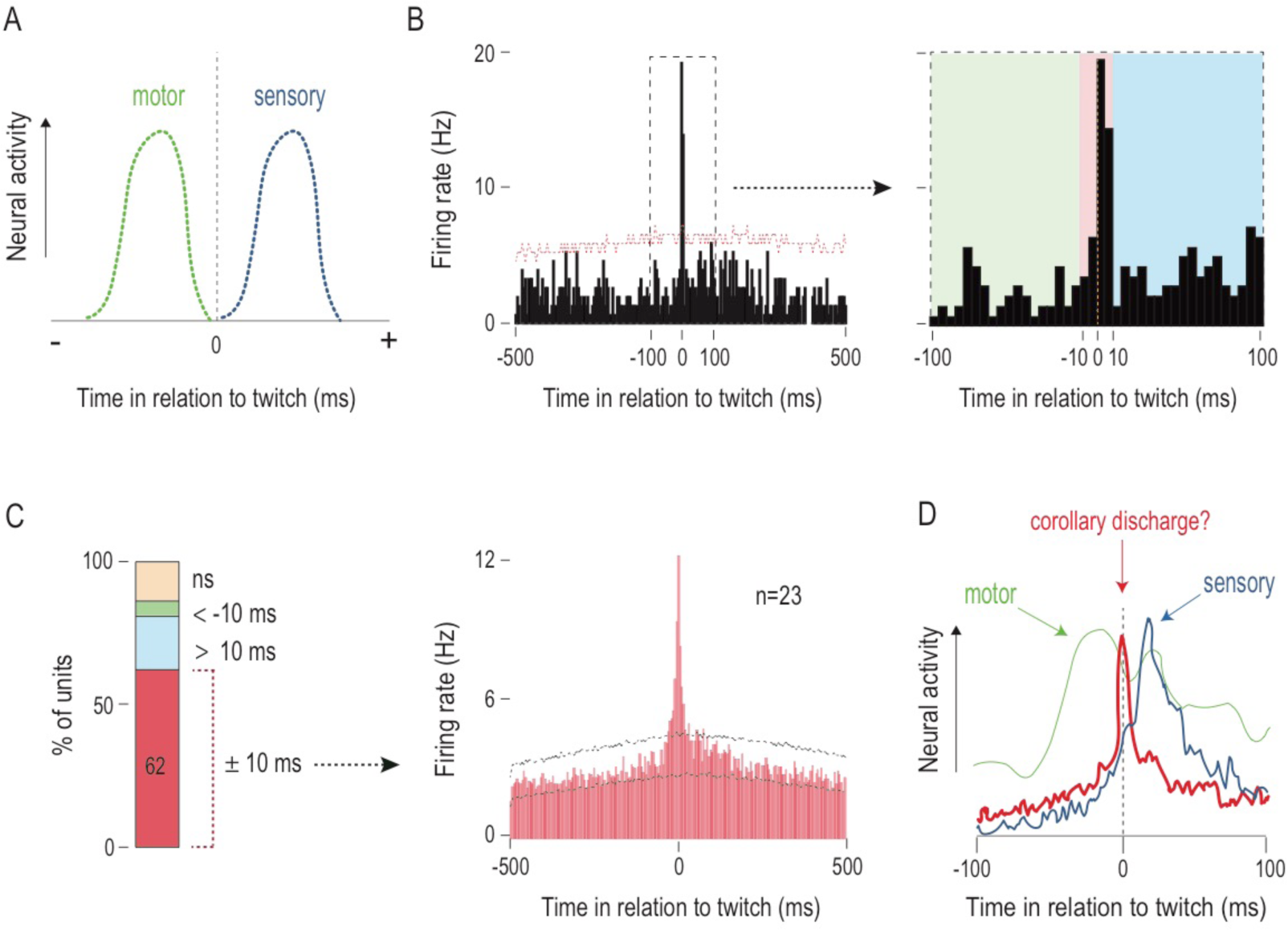
Twitches trigger sharp, short-latency olivary activity. (**A**) Perievent histograms for idealized motor and sensory units in relation to twitch onset. Motor activity precedes the onset of twitches (green line) and reafference follows the onset of twitches (blue line). Black vertical line denotes twitch onset. (**B**) Left: Perievent histogram (5-ms bins) showing sharp, short-latency activity of a representative IO unit in relation to nuchal muscle twitches. Upper confidence band (p<0.01 for each band) is indicated by red horizontal dashed line (lower confidence band is at zero). Black dashed box demarcates the ±100-ms time window around twitches. Right: The ±100-ms time window around twitches at left is shown. Green, red, and blue shaded areas denote <-10-ms, ±10-ms, and >10-ms time windows, respectively. (**C**) Left: Stacked plot showing the percentage of IO units that exhibited significant increases in firing within ±10-ms (red), >10-ms (blue), and <-10-ms (green) time windows around twitches. Right: Perievent histogram showing IO unit activity in relation to twitches for those units that were significantly active in the ±10-ms time window. Data are pooled across 23 units and triggered on 6,602 twitches. Upper and lower confidence bands (p<0.01 for each band) are indicated by black and gray horizontal dashed lines, respectively. (**D**) Comparison of IO activity in relation to twitches (red; from C) with that of a representative motor (green line; data from Del Rio-Bermudez et al., 2015) and sensory (blue line; data from Tiriac and Blumberg, 2016) structure. Black vertical dashed line denotes twitch-onset.

There are three possible explanations for these sharply peaked activation patterns observed in IO units: (a) the IO is part of the motor pathway, (b) the IO receives reafference from twitches, and (c) the IO receives CD from a motor structure that produces twitches. With respect to (a), the IO, despite being implicated in the precise timing of motor behaviors (De Zeeuw et al., 1998), is not directly involved in the generation of movements (Horn et al., 2004; Lang et al., 2017). Although it receives afferent projections from motor areas (Saint-Cyr, 1983; Saint-Cyr and Courville, 1981), there are no efferent projections from the IO to spinal motor neurons. In fact, the sole efferent projection from the IO comprises the climbing fibers that innervate cerebellar Purkinje cells (Ruigrok et al., 2014). Consequently, in adults, stimulation of the IO does not evoke or modulate movements (Gellman et al., 1985).

With respect to (b), although the IO can receive short-latency reafferent signals (Gellman et al., 1983; Sedgwick and Williams, 1967), it is unlikely that reafference can account for the short-latency peaks observed here. Consider that for the structures in which we have seen clear evidence of twitch-related reafference (e.g., ECN), we have also seen clear evidence of exafferent responses (Tiriac et al., 2014; Tiriac and Blumberg, 2016). In contrast, of the IO units that exhibited sharp peaks with a latency of +10 ms, none responded to exafferent stimulation. Moreover, of the 7 IO units that exhibited twitch-triggered responses at latencies consistent with reafferent processing (i.e., > 10 ms; Figure 2-figure supplement 1E), 3 responded to exafferent stimulation. Thus, the signature feature of the majority of IO activity—the sharp peak centered on twitch onset—is consistent with the notion that the IO receives CD associated with the production of a twitch.

### LRN neurons exhibit two kinds of twitch-related activity

Based on research in adults (Alstermark and Ekerot, 2015; Arshavsky et al., 1978), we predicted that the LRN, like the IO, would exhibit CD-related activity. Moreover, because the LRN also receives sensory inputs from the limbs (Figure 3A), we expected to see evidence of reafference in that structure. To test these two possibilities, we next recorded spontaneous LRN activity in P8 rats across sleep and wake.

**Figure 3.**
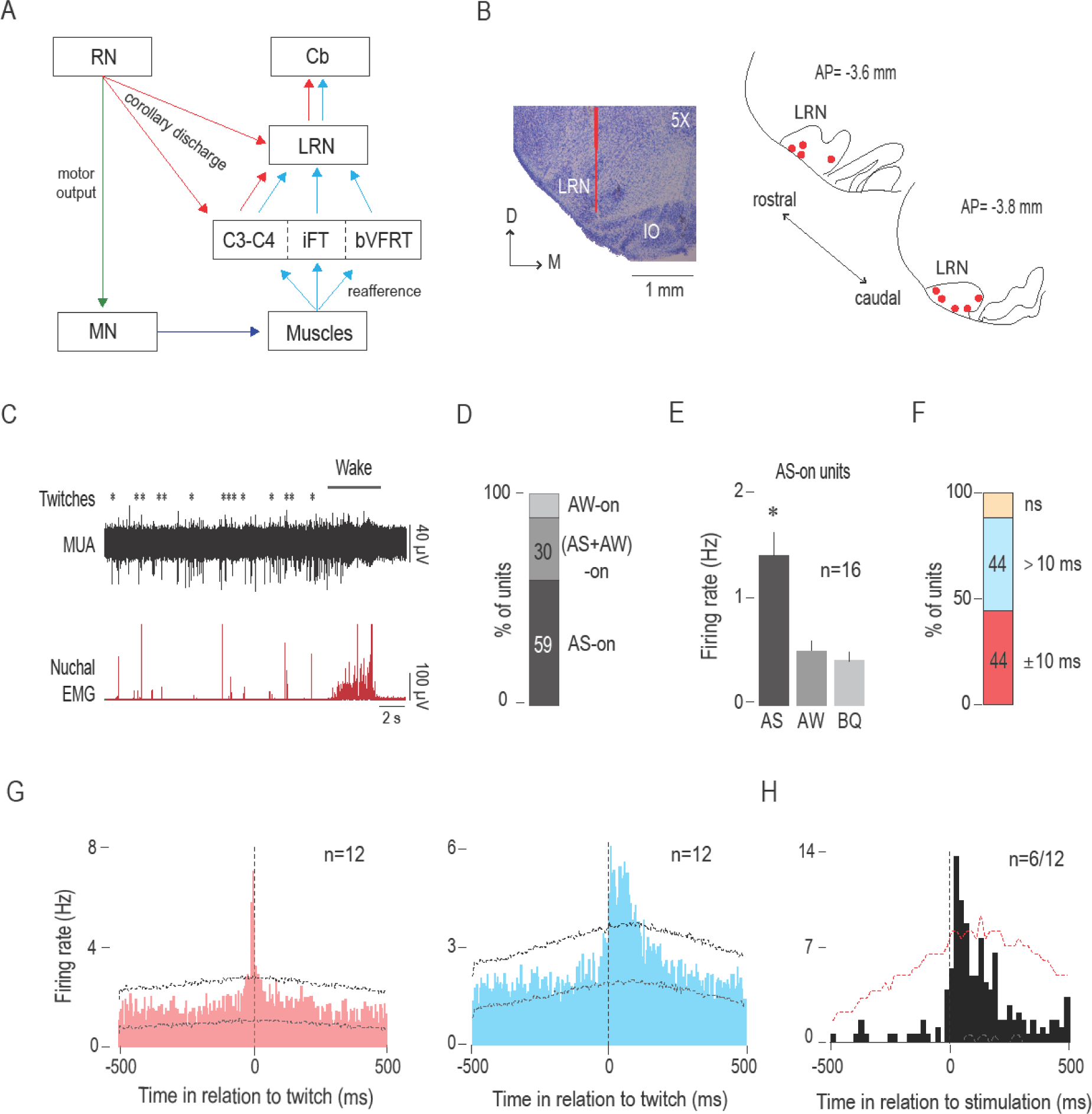
The LRN receives twitch-related corollary discharge and reafference signals. (**A**) Diagram depicting afferent and efferent connections of the LRN. Pathways conveying motor commands (green), reafference (blue), and corollary discharge (red) are shown (see Alstermark and Ekerot, 2013). (**B**) Left: Coronal section (5x) stained with cresyl violet depicting the anatomical location of the LRN in the brainstem and reconstruction of a representative electrode placement (red vertical line) in the LRN. Right: Reconstruction of electrode placements (red circles) within the LRN in two coronal sections across all P8 subjects (n=9). (**C**) Representative recording of rectified nuchal EMG activity and multiunit activity (MUA) in the LRN during spontaneous sleep-wake cycling. Asterisks denote twitches and gray horizontal bars denote wake movements as scored by the experimenter.(**D**) Stacked plot showing the percentage of AS-on, (AS+AW)-on, and AW-on units in the LRN. (**E**) Bar graphs showing mean (+SEM) firing rates of AS-on units across behavioral states. * significant difference from AW and BQ, p<0.0005. (**F**) Stacked plot depicting the percentage of units that significantly increased their firing rates ±10 ms around twitches (red) and >10 ms following twitches (blue). (**G**) Perievent histograms (5-ms bins) showing LRN unit activity in relation to twitches. Left: Data pooled across 12 units (and triggered on 3,688 twitches) that significantly increased their activity in the ±10-ms time window (red). Right: Data pooled across the 12 units (and triggered on 5264 twitches) that exhibited a significant peak with a latency of >10 ms. Black vertical dashed lines indicate twitch onset. Upper and lower confidence bands (p<0.01 for each band) are indicated by black and green horizontal dashed lines, respectively. (**H**) Perievent histogram (20-ms bins) depicting LRN unit activity (n=6 of a possible total of 12) in response to forelimb or hindlimb stimulation for those units that significantly increased their activity in the >10 ms time window (blue histogram in G). Black vertical dashed line corresponds to stimulation onset determined using EMG activity. Upper confidence band (p<0.05) is indicated by red horizontal dashed line. RN: red nucleus; MN: motor neurons; C: cervical segment; LRN: lateral reticular nucleus; Cb: cerebellum; IO: inferior olive; D: dorsal; M: medial; AP: antero-posterior in relation to lambda; AS: active sleep; AW: active wake; BQ: behavioral quiescence; ns: not significant.

We confirmed electrode placements in the LRN (n=27 units across 9 pups, 1-6 units/pup; Figure 3B). Similar to the IO, the unit activity in the LRN was phasic and restricted to periods of AS, particularly around twitches. LRN activity was sparse during BQ and was suppressed after AW onset (Figure 3C). The majority (16/27 units, 59%) of LRN units were AS-on (Figure 3D) and the mean firing rate of the AS-on units (1.4 ± 0.2 Hz) was approximately three times higher during AS than during any of the other two states (p=<0.0005; Figure 3E).

Next, we assessed the temporal relationship between unit activity and twitches by creating perievent histograms (5-ms bins, 1-s window). Regardless of state dependency, the majority of LRN units (24/27 units, 89%) showed significant increases in firing rate in response to a twitch (Figure 3F). As predicted, we observed two different neural populations the exhibited distinct patterns of twitch-triggered activity. First, we found a subpopulation of LRN units (12/27 units, 44%) that, like the majority of IO units, exhibited a sharp peak within ±10 ms of a twitch (Figure 3G, left); also, none of these LRN units responded to exafferent stimulation of the limbs (data not shown).

Second, the remaining LRN units (12/27 units, 44%) exhibited broader twitch-related activity profiles consisting of a peak in activity around twitch onset (±10 ms) and/or a peak with a latency of >10 ms (Figure 3G, right). The latter peak is what is expected from a short-latency reafferent responses (Tiriac and Blumberg, 2016; Tiriac et al., 2014). Indeed, 6 of these 12 units also responded to exafferent stimulation of either the forelimb or hindlimb with an average latency of 40 ms (Figure 3H).

### Non-overlapping regions in the mesodiencephalic junction (MDJ) project to the IO and LRN

The MDJ includes diverse structures, like the RN, that innervate spinal motor neurons (De Zeeuw et al., 1998; Saint-Cyr and Courville, 1981; Lakke, 1997; Onodera and Hicks, 2009; Zuk et al., 1983) and are therefore directly involved in the generation of movements (Fukushima, 1991; Morris et al., 2015; Onodera and Hicks, 1996; Williams et al., 2014). To determine whether MDJ neurons also project to the IO and LRN at P8, we performed retrograde tracing from each structure.

Wheat germ agglutinin (WGA) conjugated with Alexa Fluor 488 or 555 was microinjected into the IO or LRN and retrogradely labelled cell bodies were imaged using a fluorescent microscope. Retrograde tracing from the IO (n=5; Figure 4A) revealed robust labeling of cell bodies in the mesodiencephalic junction (MDJ), including the rostral interstitial nucleus of Cajal (RI), nucleus of Darkschewitsch (Dk), accessory oculomotor nuclei (MA3), and nearby diffuse areas in the MDJ. Little or no labelling was observed in the RN. In contrast, retrograde tracing from the LRN (n=4; Figure 4B) revealed robust labeling in the contralateral RN but not elsewhere in the MDJ. Finally, when the two different tracers were injected separately into the IO and LRN of the same pup (n=2; Figure 4C), we found that the LRN-projecting cell bodies were located within the RN and the IO-projecting cell bodies were located outside the RN. These findings are consistent with those in adult rats (Ruigrok et al., 2014) and suggest that the major afferent connections to the IO and LRN arise from non-overlapping structures in the MDJ at this age.

**Figure 4 and 1 supplement.**
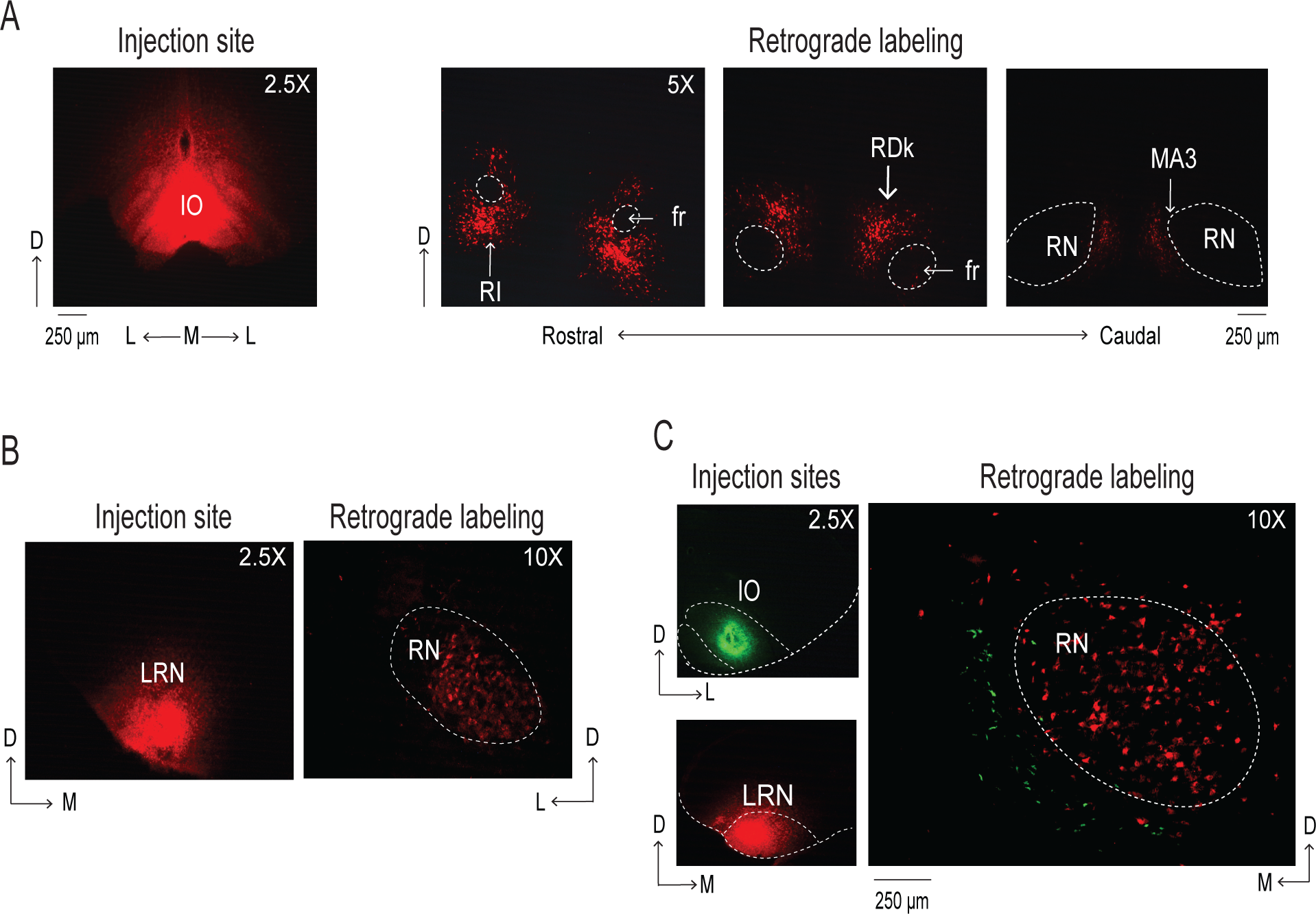
Retrograde labeling of the mesodiencephalic junction (MDJ) after infusion of WGA into the IO and LRN of P8 rats. (**A**) Left: Coronal section (2.5x) depicting WGA-555 diffusion in the IO. Right: Coronal sections (5x) depicting retrogradely labeled cell bodies in multiple MDJ nuclei. (**B**) Left: Coronal section (2.5x) depicting WGA-555 diffusion in the LRN. Right: Coronal section (10x) depicting retrogradely labeled cell bodies in the red nucleus (RN). (**C**) Left: Coronal sections (2.5x) depicting WGA-488 diffusion in the IO (top; green) and WGA-555 diffusion in the LRN (bottom; red) in the same P8 rat. Right: Coronal section (10x) depicting retrogradely labeled cell bodies in the RN (red; from LRN) and the region immediately medial to it (green; from IO). D: dorsal; L: lateral; M: medial; IO: inferior olive; RI: rostral interstitial nucleus of Cajal; fr: fasciculus retroflexus; RDk: rostral nucleus of Darkschewitsch; MA3: accessory oculomotor nuclei; LRN: lateral reticular nucleus.

### MDJ stimulation causes limb movements and c-Fos activation in the IO and LRN

To assess functional connectivity between the MDJ and the IO or LRN, we electrically stimulated the RN (n=4) and other MDJ nuclei (n=4) while monitoring forelimb and hindlimb movements in urethanized (1.5mg/g, IP) P8 rats (Figure 4—figure supplement 1A). Subsequently, we performed immunohistochemistry to determine the expression of the c-Fos protein, a marker of neural activity (Chung, 2015), in the IO and LRN.

Stimulation of several non-RN MDJ nuclei evoked non-specific movements of the ipsilateral and contralateral forelimbs and hindlimbs and also resulted in c-Fos expression primarily in the ipsilateral IO (Figure 4—figure supplement 1B). In contrast, stimulation of the RN produced only discrete contralateral forelimb movements (Figure 4—figure supplement 1A) and resulted in c-Fos expression in the contralateral LRN but not the IO (Figure 4—figure supplement 1C). These results indicate that MDJ nuclei are functionally connected to the IO and LRN at these ages.

It is possible that the c-Fos activation in the IO and LRN was due to sensory feedback arising from the stimulated movements. However, we observed little or no c-Fos expression in sensory areas like the cuneate nucleus and ECN (data not shown).

### MDJ neurons adjacent to the RN are active before and after the production of twitches

It has been established that the RN is involved in the production of twitches (Del Rio-Bermudez et al., 2015) and, as shown here, in the conveyance of CD to the LRN. Similarly, if MDJ neurons outside of the RN convey twitch-related CD to the IO, we would also expect these neurons to be involved in the production of twitches (Figure 5A). Therefore, we characterized the spontaneous activity of non-RN MDJ neurons in P8 rats during sleep and wake. We aimed to record from MDJ neurons located in regions implicated earlier as projecting to the IO and, upon stimulation, producing limb movements (see Figure 4 and Figure 4—figure supplement 1).

**Figure 5.**
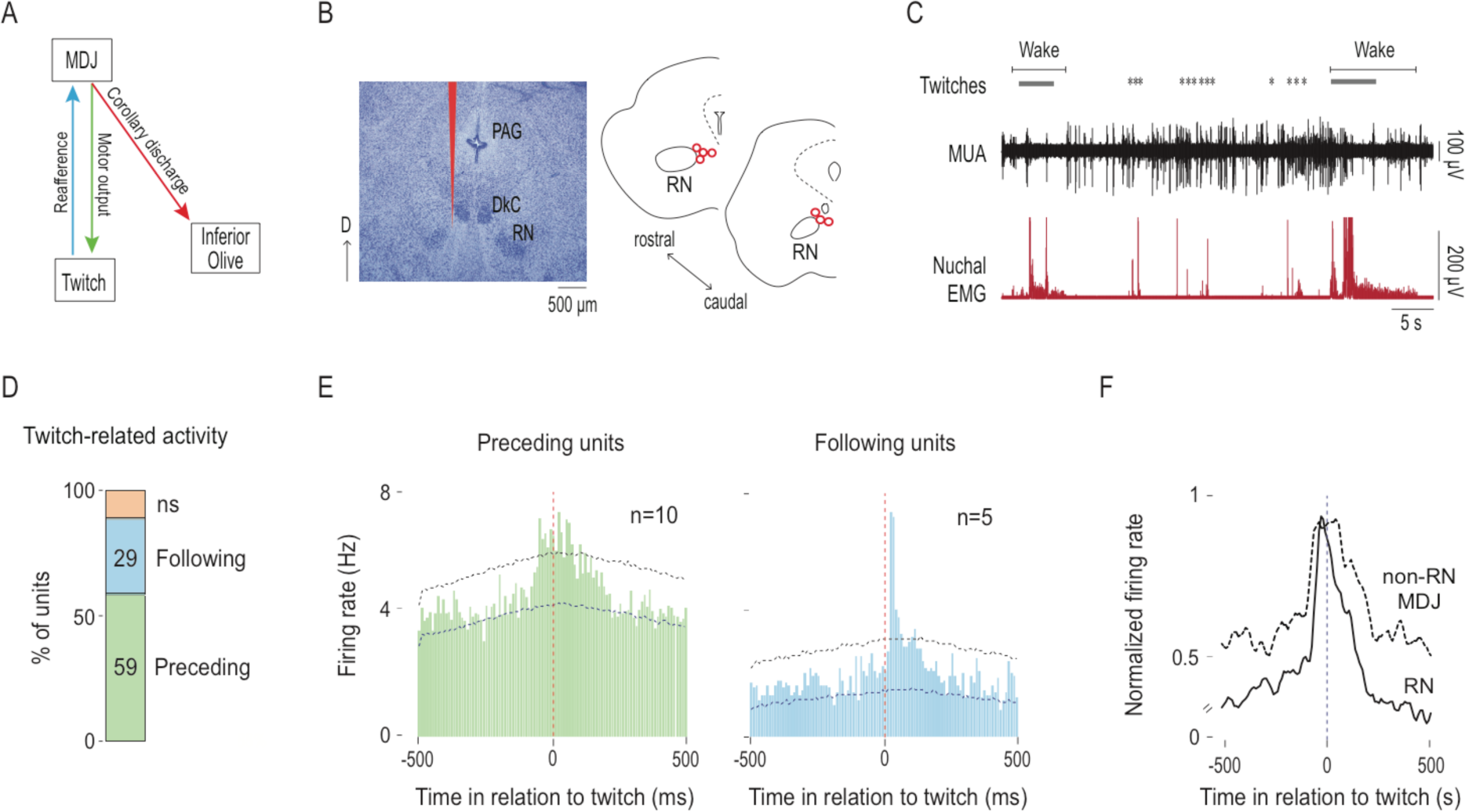
MDJ structures adjacent to the red nucleus exhibit twitch-preceding and twitch-following activity. (**A**) Diagram showing anatomical connections of the MDJ regions that lie adjacent to the red nucleus. Proposed pathways conveying motor commands (green line), reafference (blue line), and corollary discharge (red line) are shown. (**B**) Left: Reconstruction of a representative electrode placement (red vertical line; 2.5x). Right: Reconstruction of electrode placements (red circles) in the MDJ in two coronal sections across all pups (n=7). (**C**) Representative recording of rectified nuchal EMG activity and multiunit activity (MUA) in the MDJ during spontaneous sleep-wake cycling. Asterisks denote twitches and gray horizontal bars denote wake movements as scored by the experimenter. (**D**) Stacked plot depicting the percentage of twitch-preceding (motor; green) and twitch-following (sensory; blue) units in the MDJ. (**E**) Left: Perievent histogram (10-ms bins) showing activity of twitch-preceding MDJ units in relation to twitches. Data pooled across 10 units and triggered on 2877 twitches. Right: Perievent histograms (10-ms bins) showing activity of twitch-following MDJ units in relation to twitches. Data pooled across 5 units and triggered on 1382 twitches. Red vertical dashed lines correspond to twitch onset. Upper and lower confidence bands (p<0.05 for each band) are indicated by black and blue horizontal dashed lines, respectively. (**F**) Perievent histograms (10-ms bins) comparing normalized firing rate in relation to twitch onset for twitch-preceding units in the red nucleus (RN; black solid line; data from Del Rio-Bermudez et al., 2015) with that of non-RN MDJ units adjacent to the red nucleus (black dashed line; redrawn from E, left). Black vertical dashed lines correspond to twitch onsets. D: dorsal; PAG: periaqueductal gray; DkC: caudal nucleus of Darkschewitsch; ns: not significant.

Electrode placements in the MDJ immediately medial to the RN were confirmed (n=7 pups, 17 units, 1-5 units/pup; Figure 5B). The spontaneous activity of neurons in this region appeared mostly around twitches and wake movements (Figure 5C). When twitch-triggered perievent histograms (10-ms bins, 1-s window) were created, we found that the majority of the recorded units (15/17 units, 88%) showed significant twitch-dependent activity (Figure 5D).

The temporal relationship between neural activity and twitches revealed two primary subpopulations of units (Figure 5D): There were units that significantly increased their firing rates before the onset of a twitch (twitch-preceding units) and units that significantly increased their firing rates after the onset of a twitch (twitch-following units). The twitch-preceding units (10/17 units across 5 pups, 59%) showed an increase in firing rate 10-70 ms before a twitch (Figure 5E, left). Interestingly, the majority (7/10 units, 70%) of these twitch-preceding units also exhibited an increase in firing rate 10-50 ms after a twitch, indicative of reafference. The twitch-following units (5/17 units across 3 pups, 29%) showed an increase in firing rate 20-40 ms after a twitch (Figure 5E, right), suggesting they only receive reafference from a twitch. In terms of pattern and latency of twitch-triggered activity, these neurons behave similarly to those described previously in the RN (Figure 5F; Del Rio-Bermudez et al., 2015).

The twitch-preceding MDJ neurons also significantly increased their activity before the onset of wake movements (Figure 5—figure supplement 1A), consistent with that observed in the RN (Del Rio-Bermudez et al., 2015). Similarly, the IO units that exhibited twitch-related CD also significantly increased their firing rate at the onset of wake movements (Figure 5—figure supplement 1B). A similar pattern was observed in the LRN (data not shown). Therefore, the IO and LRN receive CD from the MDJ associated with both types of movements, although wake-related activity is less abundant and pronounced than twitch-related activity.

### Calcium-activated slow-potassium (SK) channels contribute to the sharp peak in IO activity

Having identified motor structures that send CD to the IO and LRN, we next sought to determine how a motor command with a broad twitch-preceding peak (see Figure 5F) is transformed into a sharp, precise peak around a twitch (see Figure 2B). We focused on the IO to address this question because of the reliably high percentage of units that exhibit twitch-related CD.

In the adult IO, SK channels prevent temporal summation of excitatory presynaptic inputs (Garden et al., 2017). SK channels are also expressed early in development (Gymnopoulos et al., 2014). Because afferent projections from the MDJ to the IO are excitatory, we hypothesized that twitch-related CD conveyed to the IO is accompanied by the opening of SK channels, thereby truncating IO activity and resulting in the observed sharp twitch-related peaks. To test this hypothesis, we blocked SK channels using apamin, an SK channel antagonist (Benington et al., 1995); apamin has been used in adult rats to block SK channels in the IO (Lang et al., 1997).

P8 rats were prepared for neurophysiological recording as described earlier. After the pup was cycling between sleep and wake, apamin (1 µM; dissolved in saline) or saline—mixed with 4% fluorogold to later identify the extent of diffusion—was microinjected at a volume of100 nl into the IO. Fifteen min after the injection (the half-life of apamin is ∼2 h; Gui et al., 2012), the microsyringe was withdrawn and was replaced with a recording electrode. Neural and EMG activity and sleep-wake behavior were then recorded for 30 min (Figure 6A).

**Figure 6 and 1 supplement.**
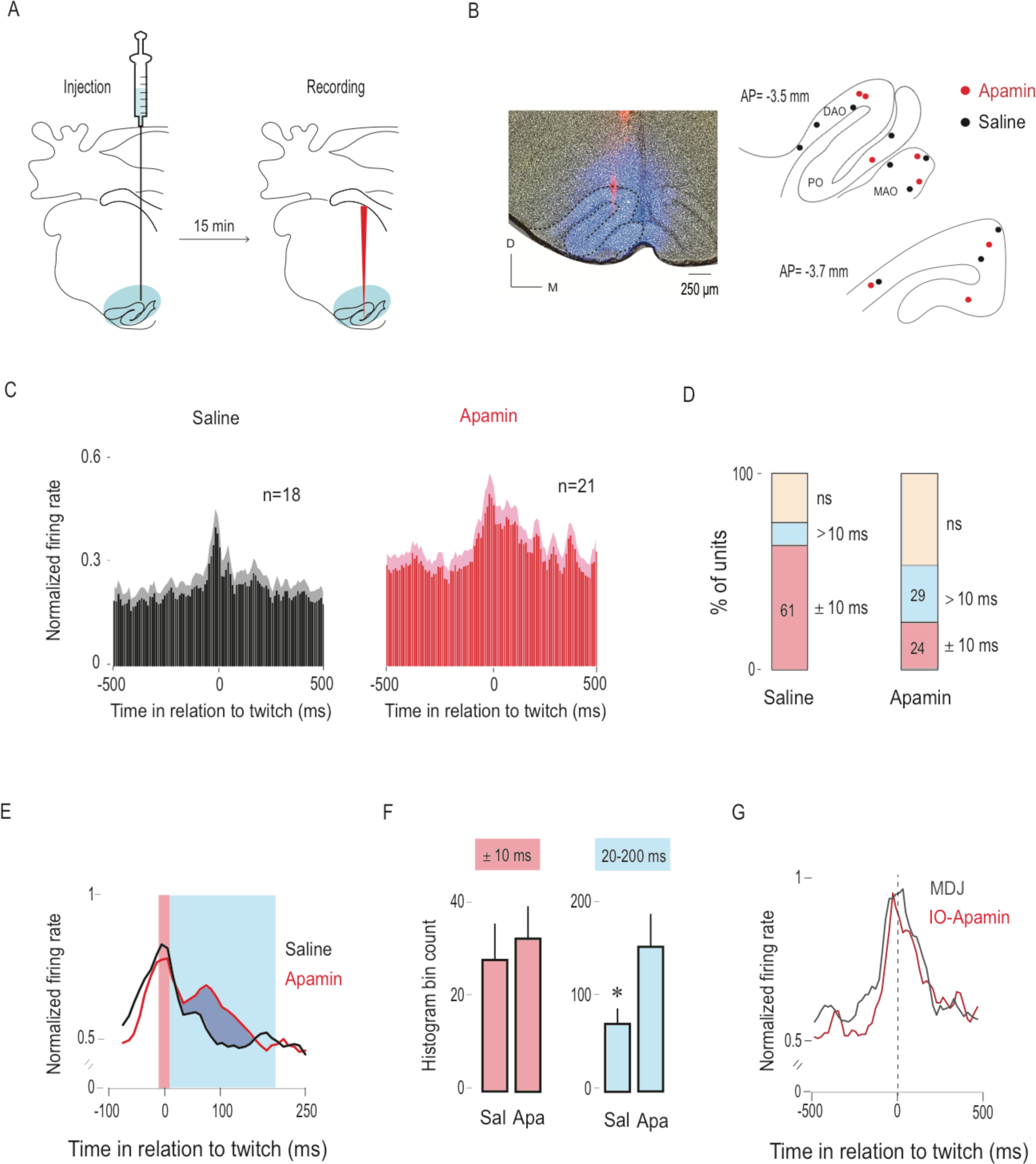
Apamin broadens the twitch-related peak in the IO. (**A**) Diagram depicting experimental design. Apamin or saline, mixed with 4% Fluorogold, was microinjected into the IO. Blue shaded area shows the extent of drug diffusion. Fifteen minutes after the injection, the microsyringe was withdrawn and a recording electrode, coated with DiI (red vertical line), was inserted into the IO. Unit activity was recorded for 30 min. (**B**) Left: Representative coronal section showing drug diffusion in the IO (blue shaded area) and placement of recording electrode (red) within the IO. Right: Reconstruction of electrode placements within the IO in two coronal sections for all pups in the saline (black dots; n=10 pups) and apamin (red dots; n=8 pups) groups. (**C**) Perievent histograms (10-ms bins) showing mean (+SEM) normalized firing rate across all units triggered on twitches in the saline (n=18) and apamin (n=21) groups. (**D**) Stacked plots showing the percentage of units with significant activity within ±10 ms around twitches (red) and >10 ms following twitches (blue) in the saline and apamin groups. (**E**) Perievent histograms (10-ms bins) showing IO unit activity in relation to twitches in the saline (black line) and apamin (red line) groups. Data are pooled across significant units only (red + blue stacks in E) in both groups (n=13 in saline and n=11 in apamin group) and smoothed (tau=10 ms). Red shaded area denotes ±10-ms time window around twitches. Blue shaded area denotes 20-200-ms time window following twitches. (**F**) Bar graphs showing the area under the curve (mean + SEM) during two time windows: the ±10-ms window around twitches (red) and the 20-200-ms window following twitches (blue) for the units in the saline (n=12) and apamin (n=11) groups. * p=0.03. (**G**) Perievent histograms comparing normalized firing rate in relation to twitch onset for twitch-preceding units in the non-RN MDJ (gray line, taken from Figure 5E) with that of IO units in the apamin group (red line). DAO: dorsal accessory olive; MAO: medial accessory olive; PO: principal olive; D: dorsal; M: medial; AP: antero-posterior in relation to bregma; sal: saline; apa: apamin; ns: not significant.

We confirmed drug or vehicle diffusion and recording sites within the IO (n=18 units across 10 pups in the saline group; n= 21 units across 8 pups in the apamin group; 1-3 units/pup; Figure 6B). There was no difference in the amount of time spent in AS (p=0.78) or in the number of twitches produced per unit time of AS between the apamin and saline groups (ps>0.15 for nuchal, contralateral and ipsilateral forelimb twitches; Figure 6—figure supplement 1A, B). As observed in the previous IO recordings, neural activity in both groups was phasic and restricted to periods of AS (Figure 6—figure supplement 1C). There was no significant difference in the overall firing rate during AS between groups (p=0.59; Figure 6—figure supplement 1D).

Perievent histograms (10-ms bins, 1-s window) were created for each individual unit in both groups (Figure 6C). As predicted, whereas twitch-triggered activity in the saline group exhibited the expected sharp peak around twitches, the activity in the apamin group was broader during the period after twitch onset.

The number of units exhibiting significant twitch-related activity did not differ between the two groups (n=13/18 in saline and n=11/21 in apamin groups; *X*^2^(1, N=39) = 1.6, p=0.2; Figure 6D). In contrast, the number of units exhibiting sharp peaks within ±10 ms of a twitch was significantly lower in the apamin group (5/21 units, 24%) than in the saline group (11/18 units, 61%; *X*^*2*^(1, N=39) = 5.6, p=0.02; Figure 6D). To illustrate the effect of apamin on twitch-related activity, we pooled the data for the significant units to create perievent histograms of IO activity. As shown in Figure 6E, the activity in the apamin group, unlike that in saline group, persisted beyond the 10-ms window after a twitch. To quantify the difference, we calculated the area under the curve for each unit during two time windows: ±10 ms around a twitch and 20-200 ms after a twitch (Figure 6F). As expected, we found no significant difference between the two groups in the ±10-ms window (*U*=56.5, Z=−0.87, p=0.4), but did find a significant difference for the 20-200-ms window, with the apamin group being significantly larger (*U*=30.5, Z=−2.2, p=0.03). In fact, the pattern of twitch-triggered neural activity in the apamin group was similar to that recorded in the MDJ (Figure 6G). Based on these results, we conclude that SK channels are involved in sharpening the CD signal arriving from the MDJ.

## Discussion

Several criteria have been proposed for identifying CD signals (Poulet and Hedwig, 2007; Sommer and Wurtz, 2008). First, a CD should originate in a structure that is demonstrably involved in the production of movement; as shown here, the twitch-related activity in the IO and LRN originates from several independent motor structures in the MDJ that are involved in the production of twitches and wake movements (Del Rio-Bermudez et al., 2015). Second, areas receiving CD should themselves play no direct role in the production of movement; this is clearly true of the IO and LRN (Gellman et al., 1985; Ruigrok et al., 2014). Third, neurons receiving CD should increase their activity at the onset of a movement; as shown here, the activity of IO and LRN neurons occurs precisely at the onset of twitches, exhibiting a temporal profile that clearly distinguishes it from twitch-preceding activity in the MDJ nuclei and twitch-following activity in the ECN. Thus, the twitch-related activity in the IO and LRN satisfies the key criteria of CD. Below we discuss the implications of this finding and its significance for sensorimotor development.

### Neurophysiological identification of CD signals in behaving animals

Neural pathways conveying CD have been delineated in a diverse array of species (Dale and Cullen, 2017; Davis et al., 1973; Fee et al., 1997; Schneider et al., 2014; Sommer and Wurtz, 2002; Yang et al., 2008). Neural recordings of the CD signal itself, however, have mostly been performed in non-mammalian species, including crickets, sea slugs, crayfish, tadpoles, and electric fish (Evans et al., 2003; Kirk and Wine, 1984; Li et al., 2004; Poulet and Hedwig, 2006; Requarth and Sawtell, 2014). The relatively small and simple nervous systems of these species have allowed for the isolation of neurons that carry or receive CD signals and identify their relationship to behavior. In contrast, CD signals have thus far only been recorded in the mediodorsal thalamus of non-human primates during eye movements (Sommer and Wurtz, 2004) and the auditory cortex of mice (Schneider et al., 2014).

The current findings provide the first direct neurophysiological evidence of CD in a developing mammal. Moreover, this is the first direct evidence of CD in the IO and LRN, consistent with what has been proposed for these two structures (Alstermark and Ekerot, 2013; Arshavsky et al., 1978; De Zeeuw et al., 1998; Devor, 2002). Also, with this discovery of a unique neural CD signature—comprising a short-latency onset and sharp activity peak—we have a clear template to guide future neurophysiological investigations of CD signals in other species and neural systems across the lifespan.

### A neural mechanism for sharpening the CD signal

As mentioned above, one of the signature features of the twitch-related CD signal is the sharp peak. This is surprising because, as shown here and in a previous study (Del Rio-Bermudez et al., 2015), twitch-related motor activity in MDJ neurons exhibits broad peaks (see Figure 5F). How does a broad presynaptic signal in the MDJ get converted into a sharp postsynaptic response in the IO and LRN (see Figures 2C, 3G)? To answer this question, we focused on the IO because, compared with the LRN, a much higher proportion of its neurons exhibited sharp peaks.

There are a few possible candidate mechanisms. For example, in cortical pyramidal neurons, interactions between excitatory and inhibitory inputs can sharpen a neuron’s activity profile (Kremkow et al., 2010). A similar mechanism is unlikely to operate in the IO for several reasons. First, inhibitory interneurons are sparse in that structure (<0.1%; Nelson and Mugnaini, 1988). Second, although the IO receives its predominant inhibitory input from the deep cerebellar nuclei (DCN; de Zeeuw et al., 1988), DCN activity occurs ∼40 ms after a twitch (Del Rio-Bermudez et al., 2016). Moreover, in pilot experiments, we found that pharmacological inactivation of the DCN had no effect on IO activity at P8, consistent with a previously published report (Nicholson and Freeman, 2003).

Consequently, we hypothesized that inhibition in the IO is mediated by SK channels. In the IO of adult rats, these channels prevent summation of excitatory inputs (Garden et al., 2017). Here, using pharmacological inactivation, we demonstrate that SK channels are responsible for sharpening the olivary CD signal. A similar mechanism could be functional in the LRN as SK channels are also expressed in that structure in adult rats (Xu et al., 2013).

### Differential actions of CD signals at precerebellar nuclei

In a previous study (Tiriac and Blumberg, 2016), it was demonstrated that wake-related reafference is blocked within the ECN, an effect that we attributed to modulation by a wake-related CD signal. That CD-mediated blockade was lifted during twitching, thereby allowing twitch-related reafference to be conveyed to downstream motor structures, including the cerebellum. In contrast, focusing here on the IO and LRN, we found that the twitch-related CD signals themselves—not reafference—are conveyed to the cerebellum (Figure 7A). Therefore, within this broader context, we see that CD accompanies sleep and wake behavior in infants, but its effects are not monolithic: It can modulate the action of a comparator to gate reafference (as in the ECN) or be transmitted sequentially to multiple downstream structures (as in the IO or LRN → cerebellum). Such diverse effects of CD have been described elsewhere (Crapse and Sommer, 2008).

**Figure 7.**
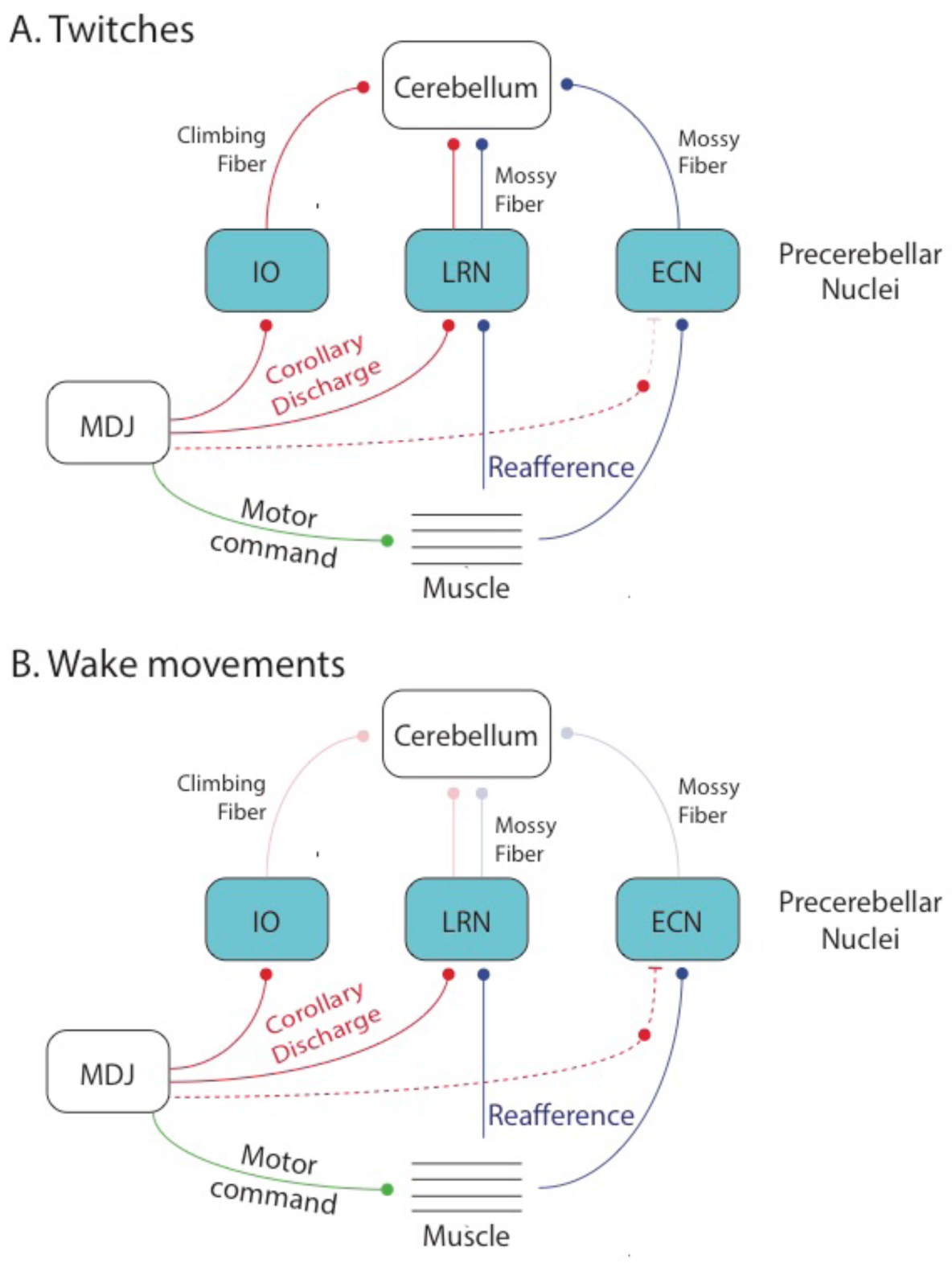
Summary diagrams depicting flow of neural activity in the cerebellar system in association with twitches and wake movements in week-old rats. (**A**) Flow of twitch-related CD from the MDJ to the IO and LRN and reafference from the limbs to the LRN and ECN. Both CD and reafference converge in the cerebellum via climbing and mossy fibers. Dotted lines denote hypothesized functional connections. (**B**) Activity is less pronounced or actively suppressed during wake movements in these three precerebellar nuclei, thereby reducing input to the cerebellum.

Although the three precerebellar nuclei—IO, LRN, and ECN—process CD and reafference differently, the common denominator of all this activity is the inundation of the developing cerebellum with twitch-related information (Figure 7A). In contrast, wake-related activity has been shown to be blocked at the level of the ECN (Tiriac and Blumberg, 2016), thereby accounting in part for the relatively low levels of activity in the cerebellum at this age (Sokoloff et al., 2015a; Sokoloff et al., 2015b). In addition, we found here that the IO and LRN are also considerably less active during periods of wake. Although wake movements were occasionally accompanied by bursts of IO and LRN activity, this activity was much less pronounced than that observed at the onset of twitching. These weaker activation patterns could simply be due to the fact that wake movements, compared to twitches, are fewer in number at this age. For example, for a typical limb muscle in a P8 rat over a 30-min period, we find that there are ∼300 twitches but only ∼30 wake movements. Consequently, when compared with twitches, there are many fewer opportunities for wake movements to activate downstream cerebellar structures (Figure 7B).

### Functional implications of CD and reafference

The developing sensorimotor system receives substantial sensory input from self-generated twitches and from external stimulation arising from the mother and littermates. It has been suggested that the infant brain does not distinguish between these two sources of input and that twitch-related reafference serves merely as a “proxy” for exafferent stimulation (Akhmetshina et al., 2016; McVea et al., 2016). This suggestion rests in part on the observation that both forms of stimulation, despite their very different origins, trigger similar patterns of cortical activity (Akhmetshina et al., 2016; Tiriac et al., 2012; Yang et al., 2013). Thus, with our finding that CD accompanies the production of twitches, it is now clear that there exists a mechanism with which the infant brain can distinguish self-generated from other-generated movements; the ability to make this distinction is thought to rely in part on the cerebellum (Blakemore et al., 2000; Wolpert et al., 1998).

There are a number of ways in which twitch-related CD could contribute to cerebellar development and function. For example, in the adult cerebellum, CD and reafference are known to converge via climbing and mossy fibers (Blakemore et al., 2001; Huang et al., 2013; van Kan et al., 1993; Wolpert et al., 1998). In this way, it is thought that the cerebellum instantiates a forward model that receives sensory predictions and computes prediction errors (by comparing CD with reafference) in order to facilitate motor learning (Blakemore et al., 2000; Brooks et al., 2015; Requarth and Sawtell, 2014; Wolpert et al., 1998).

Twitches could contribute to the process by which forward models are instantiated and updated, especially in the context of a rapidly growing body. To appreciate this possibility, consider this description of cerebellar function: “After much trial and error during infancy and throughout life, the cerebellum learns to associate actual movements with intended movements. Many of our motor memories are movements that we have repeated millions or billions of times…” (p. 538, Mason, 2011). In that context, the millions of twitches produced in early infancy could be a critical source of repeated convergent input to the developing cerebellum. This convergence, illustrated in Figure 7, would provide the developing cerebellum with abundant opportunities to align prediction and feedback signals in a topographically organized fashion.

Cerebellar circuitry undergoes substantial development over the first three postnatal weeks in rats (Altman, 1972a; 1972b; 1972c; Shimono et al., 1976; Wang and Zoghbi, 2001). Many of these developmental processes depend heavily on neural activity, including climbing fiber synapse elimination and translocation at Purkinje cells (Andjus et al., 2003; Kakizawa et al., 2000; Kano and Hashimoto, 2012; Watanabe and Kano, 2011). With respect to synapse elimination, beginning around P8, the initial multiple innervation of Purkinje cells by climbing fibers begins to shift toward singly innervated cells in the second postnatal week as one climbing fiber is selectively strengthened over others. Importantly, spike timing-dependent plasticity (STDP) has been implicated in this process (Kawamura et al., 2013); STDP depends on the repetitive and sequential firing of pre-and post-synaptic cells within a short and precise time window (Feldman, 2012; Kawamura et al., 2013; Sgritta et al., 2017). The present findings in precerebellar nuclei, in which twitch-related CD reliably preceded reafference by approximately 10-30 ms, are consistent with twitches playing a role in cerebellar development via STDP. Moreover, recording from Purkinje cells at P8, we previously found that complex and simple spikes were highly likely to occur within 0-50 ms after twitches (Sokoloff et al., 2015a).

### Implications for neurodevelopmental disorders

Disruption of cerebellar function during sensitive periods of development can have negative cascading effects on cerebello-cortical communication and ultimately on associated sensorimotor and cognitive processes, as observed in autism spectrum disorder (Diamond, 2000; Wang et al., 2014). There are many potential causes of early cerebellar dysfunction, including prenatal and postnatal exposure to environmental stressors (Wang et al., 2014). One such stressor could be sleep deprivation or restriction, especially during early infancy when sleep is the predominant state (Jouvet-Mounier et al., 1970; Roffwarg et al., 1966). As demonstrated here and in previous studies (Sokoloff et al., 2015a; Sokoloff et al., 2015b), AS provides an important context for cerebellar activity in early development. Therefore, perturbation of sleep could deprive the cerebellum and other structures of critical sensorimotor activity during sensitive periods of development.

Accumulating evidence also suggests that CD-related processing is dysfunctional in patients with schizophrenia. Specifically, failure to disambiguate “self-generated” from “other-generated” sensory input may underlie hallucinations and delusions of control (Feinberg and Guazzelli, 1999; Ford et al., 2008). If twitches help to instruct the developing brain to distinguish self from other, disruption of sleep and sleep-related sensorimotor processing may have later-emerging negative consequences for the processing of CD.

### Conclusion

It has been argued that the discreteness of twitches makes them ideally suited to provide high-fidelity sensory information at ages when activity-dependent development is so important for the developing nervous system (Blumberg et al., 2013; Tiriac et al., 2015). The present results go further to suggest that the convergence of twitch-related CD and reafference associated with millions of twitches over the early developmental period provides ample opportunity for assimilating growing limbs into the infant’s emerging body schema (Blumberg and Dooley, 2017).

## Materials and Methods

All experiments were carried out in accordance with the National Institutes of Health Guide for the Care and Use of Laboratory animals (NIH Publication No. 80-23). Experiments were also approved by the Institutional Animal Care and Use Committee (IACUC) of the University of Iowa.

### Subjects

Male and female Sprague-Dawley Norway rats *(Rattus norvegicus)* at postnatal day (P) 7-9 (hereafter P8; n=68) from 60 litters were used for the study. All litters were culled to eight pups by P3. Mothers and litters were housed and raised in standard laboratory cages (48 × 20 × 26 cm). Food and water were available ad libitum. The animals were maintained on a 12-h light-dark cycle with lights on at 0700 h. Littermates were never assigned to the same experimental group.

### Surgery

A pup with a visible milk band was removed from the home cage. Under isoflurane (3-5%) anesthesia, bipolar hook electrodes (50 µm diameter, California Fine Wire, Grover Beach, CA) were inserted into the nuchal, forelimb, and hindlimb muscles for electromyography (EMG) and secured with collodion. A stainless steel ground wire was secured transdermally on the back. A custom-built head-fix device was then secured to the exposed skull with cyanoacrylate adhesive (Blumberg et al., 2015). The local anesthetic, Bupivicaine (0.25%) was applied topically to the site of incision and some subjects were also injected subcutaneously with the analgesic agent carprofen (0.005 mg/g). The pup was lightly wrapped in gauze and allowed to recover in a humidified, temperature-controlled (35 °C) incubator for at least one hour. After recovery, the pup was briefly (<15 min) re-anesthetized with isoflurane (2-3%) and secured in a stereotaxic apparatus. A hole was drilled in the skull for insertion of the recording electrode into the inferior olive (IO; coordinates: AP=3.4-3.6 mm caudal to lambda; ML=0-1.2 mm), the lateral reticular nucleus (LRN; coordinates: AP=3.5-3.7 mm caudal to lambda; ML=1.5-1.8 mm), or midbrain nuclei near the red nucleus (RN) within the mesodiencephalic junction (MDJ; coordinates: AP=4.7-4.9 mm caudal to bregma; ML=0.2-0.5 mm). Two additional holes were drilled over the frontal or parietal cortices for subsequent insertion of the ground wire and a thermocouple (Omega Engineering, Stamford, CT) to measure brain temperature. In 15 pups in which no neurophysiological recordings were performed, only one additional hole was drilled for insertion of the thermocouple. After surgery, the pup was transferred to the recording chamber.

### Electrophysiological recordings

The head-fix device was secured to the stereotaxic apparatus housed within the recording chamber and the pup was positioned with its body prone on a narrow platform with limbs dangling freely on both sides (Blumberg et al., 2015). Care was taken to regulate air temperature and humidity such that the pup’s brain temperature was maintained at 36-37 °C. Adequate time (1-2 h) was allowed for the pup to acclimate to the recording environment and testing began only when it started cycling normally between sleep and wake. Pups rarely exhibited abnormal behavior or any signs of discomfort or distress; when they did, the experiment was terminated. The bipolar EMG electrodes were connected to a differential amplifier (A-M Systems, Carlsborg, WA; amplification: 10,000x; filter setting: 300-5000 Hz). A ground wire (Ag/AgCl, 0.25 mm diameter, Medwire, Mt. Vernon, NY) was inserted into the frontal or parietal cortex contralateral to the recording site and a thermocouple was inserted into the frontal or parietal cortex ipsilateral to the recording site. Neurophysiological recordings were performed using a 16-channel silicon electrode or a 4-channel linear probe (A1×16-10mm-100-177; A1×16-8mm-100-177; Q1×4-10mm-50-177, NeuroNexus, Ann Arbor, MI), connected to a data acquisition system (Tucker-Davis Technologies, Alachua, FL) that amplified (10,000x) and filtered (500-5000 Hz) the neural signals. A digital interface and Spike2 software (Cambridge Electronic Design, Cambridge, UK) were used to acquire EMG and neurophysiological signals at 1 kHz and at least 12.5 kHz, respectively.

A micromanipulator (FHC, Bowdoinham, ME) was used to lower the electrode into the brain (DV; IO: 5.5-6.2 mm, LRN: 5-5.8 mm, midbrain structures: 4.5-4.9 mm) until action potentials were detected. Recording began at least 10 min after multiunit activity (MUA) was detected. Before insertion, the electrode was dipped in fluorescent DiI (Life Technologies, Grand Island, NY) for later identification of the recording sites. Recording of MUA and EMG activity continued for 30 min as the pup cycled freely between sleep and wake (in two pups, activity was recorded for only 15 min). The experimenter, blind to the electrophysiological record, scored the pup’s sleep and wake behaviors, as described previously (Karlsson et al., 2005).

At the end of the recording session, the experimenter assessed evoked neural responses to exafferent stimulation of the limbs. Forelimbs and hindlimbs were gently stimulated using a paint brush. When evoked responses were observed in at least one of the recording channels, stimulation was repeated 20-30 times at intervals of at least 5 s. Each stimulus event was marked using a key press.

### Histology

At the end of all recording sessions, pups were anesthetized with sodium pentobarbital (1.5 mg/g IP) or ketamine/xylazine (0.02 mg/g IP) and perfused transcardially with phosphate-buffered saline and 4% formaldehyde. Brains were sectioned coronally at 80µm using a freezing microtome (Leica Microsystems, Buffalo Grove, IL). Recording sites were determined by examining DiI tracks, before and after staining with cresyl violet, using a fluorescent microscope (Leica Microsystems, Buffalo Grove, IL).

### Retrograde tracing

Retrograde tracing was performed at P8 (n=9) using wheat germ agglutinin (WGA) conjugated to Alexa Fluor 555 or 488 (Invitrogen Life Technologies, Carlsbad, CA). WGA-555 was injected into the IO in 3 pups and into the LRN in 2 pups. In the remaining 2 pups, dual tracing was performed by injecting WGA-488 into the IO and WGA-555 into the LRN. To perform these injections, a pup was anesthetized with 2-5% isoflurane and secured in a stereotaxic apparatus. A 0.5 µl microsyringe (Hamilton, Reno, NV) was lowered stereotaxically into the IO or LRN and 0.01-0.02 µl of 2% WGA-555 or WGA-488 (dissolved in 0.9% saline) was injected over 1 min. After a 15-min post-infusion period, the microsyringe was withdrawn and the incision was closed using Vetbond (3M, Maplewood, MN). The pup was returned to its home cage and perfused 24 h later as described above. Brains were sectioned coronally at 50 µm. Every other section was kept for Nissl staining for verification of the injection sites and areas that show retrograde labeling. Retrogradely labeled cell bodies were imaged using a fluorescent microscope (DFC300FX, Leica, Buffalo Grove, IL)

### Stimulation of MDJ structures

In urethanized (1.5 mg/g) head-fixed P8 rats (n=8), a parylene-coated tungsten stimulating electrode (World Precision Instruments, Inc., Sarasota, FL) was lowered into the MDJ nuclei most strongly implicated by retrograde tracing. The nuclei were electrically stimulated to produce discrete movements of the forelimbs and/or hindlimbs. Trains of pulses (pulse duration: 0.2-0.4 ms; pulse frequency: 300 Hz; train width: 45 ms; Williams et al., 2014) were delivered every 5 s for 60 min. The current was adjusted (300-900 μA) as needed to ensure that stimulation continued to reliably produce movement. Ninety min after the last stimulation, the pup wa sacrificed and the brain was prepared for c-Fos immunohistochemistry.

### Immunohistochemistry for c-Fos expression

Brains were sliced in 50 μm sections and every other section was kept for Nissl staining for verification of the stimulation sites and visualization of c-Fos expression, respectively. Primary antibody against c-Fos (anti-c-Fos rabbit polyclonal IgG; Santa Cruz Biotechnology) was diluted 1:1000 in a universal blocking serum (2% bovine serum albumin;1% triton;0.02% sodium azide) and applied to the sections. Sections were coverslipped and left to incubate for 48 h at 4°C. After incubation of the primary antibody and a series of washes in PBS, a secondary antibody (Alexa Fluor 488 donkey anti-rabbit IgG; Life Technologies, Grand Island, NY; 1:500 in PBS) was applied to the section and incubate for 90 min at room temperature. The slides were coverslipped using Fluoro-Gel (Electron Microscopy Sciences (Hatfield, PA) and expression of c-Fos was examined using a fluorescent microscope (DFC300FX or DM6B, Leica, Buffalo Grove, IL).

### Intra-IO injection of apamin

In 18 P8 rats, pups were prepared for electrophysiological recording as described above and transferred to the recording rig. Once a pup started cycling between sleep and wake, a 0.5 µl microsyringe was lowered stereotaxically into the IO and 100 nl of apamin. (Abcam, Cambridge, MA; 1 µM, dissolved in 0.9% saline, n=8) or saline (n=10) was injected over 1 min. During preparation of the drug or vehicle, fluorogold (4%, Fluorochrome, Denver, CO) was added to the solutions for subsequent assessment of the extent of drug diffusion. After a 15-min period to allow for diffusion, the microsyringe was withdrawn and a recording electrode was lowered in its place into the IO and activity was recorded for 30 min. At the end of the experiment, the pup was sacrificed and its brain was prepared for histology as described above.

### Data analysis

#### Spike sorting

As described previously (Mukherjee et al., 2017; Sokoloff et al., 2015a), action potentials (signal-to-noise ≥ 2:1) were sorted from MUA records using template matching and principal component analysis in Spike2 (Cambridge Electronic Design). Waveforms exceeding 3.5 SD from the mean of a given template were excluded from analysis.

#### Identification of behavioral states

EMG activity and behavioral scoring were used to identify behavioral state (Blumberg et al., 2015). To establish an EMG threshold for distinguishing sleep from wake, EMG signals were rectified and smoothed (tau = 0.001 s). The mean amplitude of high muscle tone and atonia were calculated from five representative 1-s segments and the midpoint between the two was used to establish the threshold for defining periods of wake (defined as muscle tone being above the threshold for at least 1 s) and sleep (defined as muscle tone being below the threshold for at least 1 s). Active wake (AW) was identified by high-amplitude limb movements (e.g., stepping, stretching) against a background of high muscle tone and was confirmed using behavioral scoring. The onset of a wake movement was defined on the basis of EMG amplitude surpassing the established threshold. Active sleep (AS) was characterized by the presence of myoclonic twitches of the limbs against a background of muscle atonia. Twitches were identified as sharp EMG events that exceeded by ≥ 3x the mean EMG baseline during atonia; twitches were also confirmed by behavioral scoring (Seelke and Blumberg, 2010). Additionally, behavioral quiescence (BQ) was characterized as periods of low muscle tone interposed between AW and AS.

#### State-dependent neural activity

For each unit, average firing rate across all behavioral states was determined. Bouts of AS, AW, and BQ were excluded when firing rates exceeded 3 SD of the firing rate for that behavioral state; this happened rarely (0-2 per unit). Next, pairwise comparison of firing rates across states was performed using the Wilcoxon matched-pairs signed-ranks test (SPSS; IBM, Armonk, NY). Units were categorized as AS-on (AS > AW ≥ BQ), AW-on (AW > AS ≥ BQ), AS+AW-on (AS = AW > BQ) or state-independent (AS = AW = BQ). Firing rates of all AS-on units across behavioral states were further compared using the Wilcoxon matched-pairs signed-ranks test.

#### Twitch-triggered neural activity

To determine the relationship between unit activity and twitching, we triggered unit activity on twitch onsets and generated perievent histograms over a 1-s window using 5-or 10-ms bins. We performed these analyses on each individual unit using twitches from nuchal, forelimb, and hindlimb muscles. We tested statistical significance by jittering twitch events 1000 times over a 500-ms window using PatternJitter (Amarasingham et al., 2012; Harrison and Geman, 2009). Then using a custom-written Matlab program (MathWorks, Natick, MA), we generated upper and lower confidence bands (p<0.05 or 0.01 for each confidence band) using a method that corrects for multiple comparisons (Amarasingham et al., 2012). For each unit, after histograms were separately constructed for nuchal, forelimb, or hindlimb twitches, we identified and activity that was significant in response to a twitch. When more than one muscle yielded a significant change in neural activity, we further analyzed the data only for the muscle that showed the strongest relationship (determined by the highest firing rate) between twitches and unit activity. We then pooled these data to create perievent histograms composed of significant units and performed jitter analyses on the pooled data.

#### Wake-triggered neural activity

To determine the relationship between neural activity and wake movements, we triggered unit activity on wake-movement onset and created perievent histograms on the pooled data (20-ms bins, 1-s window). We then performed jitter analysis as described above.

#### Evoked response to exafferent stimulation

. We identified MUA in which evoked responses were observed and then sorted the units. Those units were then pooled and triggered on stimulus onset (determined using EMG artifact) to create perievent histograms. The jitter analysis was performed on the pooled data, as described above.

#### Intra-olivary injection of apamin

First, we identified if apamin affected sleep-wake behavior. We assessed the amount of time spent in AS and the number of twitches per min of AS in each pup. Differences across groups were tested using the Mann-Whitney *U* test. Next, we determined if apamin altered the overall firing rate. We calculated the firing rate of each unit during AS and compared that across groups using the Mann-Whitney *U* test. One value exceeding 3 SD was excluded as an outlier.

We then assessed whether apamin altered the shape of twitch-triggered perievent histogram. First, we created perievent histograms (10-ms bins, 1-s window) for each unit as described above. For each unit, firing rate was normalized to the peak firing rate and the average normalized firing rate across all units in each group was calculated. Perievent histograms were then created with the average (+SEM) normalized firing rates triggered on twitches for each group. Next, we assessed how apamin altered the pattern of twitch-triggered activity of individual units. To do that, we identified significant units by performing jitter analysis on individual units as described above. We counted the percentage of units that showed precise peak within ±10 ms around a twitch and compared that across groups using a Chi-squared test. Finally, we pooled significant units in each group and pooled them to create perievent histograms consisting of significant units only. To assess the difference in the shape of perievent histograms, we calculated the area under the curve by adding the histogram counts within a particular time window and compared that across groups using the Mann-Whitney *U* test. One value exceeding 3 SD was excluded as an outlier.

Unless otherwise stated, alpha was set at 0.05.

## Figures and Legends

**Figure 2—figure supplement 1.**
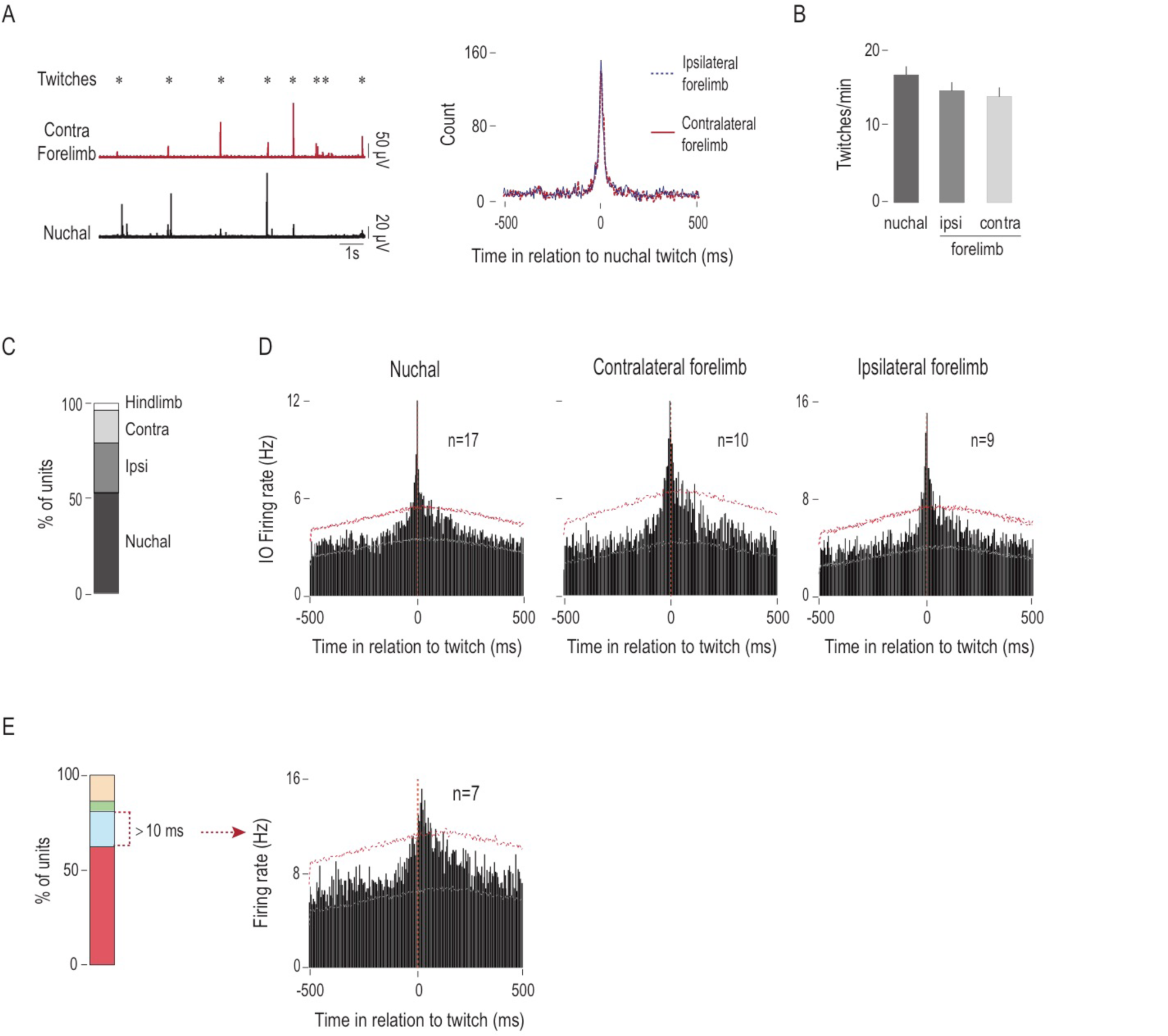
IO units respond predominantly to nuchal and forelimb twitches. (**A**) Left: Representative recording of nuchal and contralateral forelimb EMG activity showing co-occurrence of twitches in those muscles. Asterisks denote experimenter-scored twitches. Right: Cross-correlogram (5-ms window) depicting co-occurrence of nuchal muscle twitches with contralateral (red line) and ipsilateral (blue dotted line) forelimb twitches. (**B**) Bar graphs depicting mean (+SEM) rates of twitching in nuchal muscle and ipsilateral and contralateral forelimb muscles. Data are pooled across 15 pups (**C**) Stacked plot showing the percentage of IO units that exhibited sharp, zero-latency activity in relation to nuchal, ipsilateral forelimb, contralateral forelimb, and hindlimb twitches. (**D**) Perievent histograms showing IO unit activity in relation to nuchal muscle and contralateral and ipsilateral forelimb twitches. Data are pooled across 17, 10, and 9 units, respectively. Upper and lower confidence bands (p<0.01 for each band) are indicated by red and gray horizontal dashed lines, respectively. ipsi: ipsilateral; contra: contralateral. (**E**) Left: Stacked plot showing percentages of IO units that exhibited significant increase in firing in >10-ms (blue stack) time windows around twitches. Right: Perievent histograms showing IO unit activity in relation to twitches. Data pooled across 7 units. Ipsi=ipsilateral; contra=contralateral; ns=not significant.

**Figure 4—figure supplement 1.**
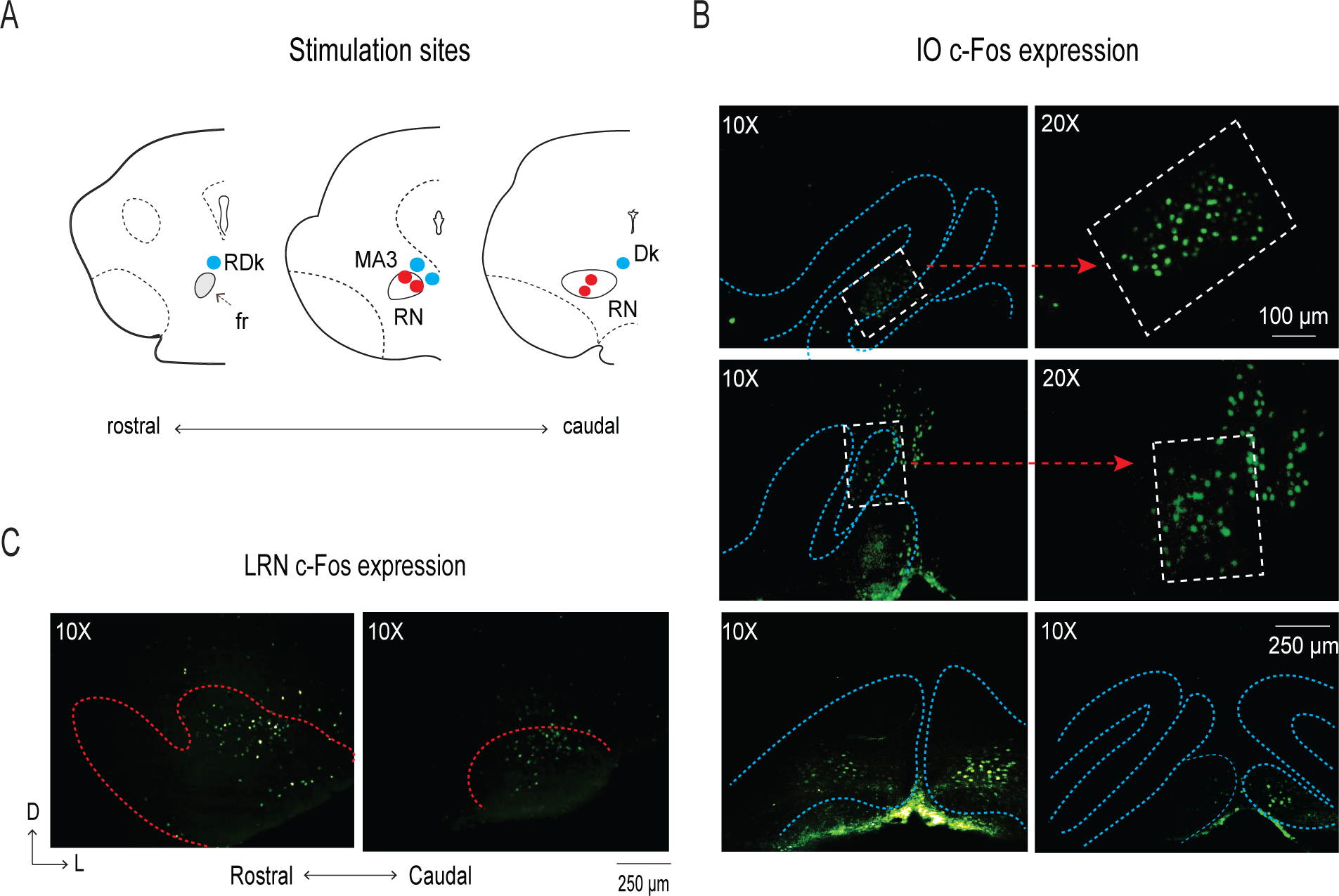
(**A**) Reconstruction of stimulation sites inside (red dots) and adjacent to (blue dots) the red nucleus (RN) in three coronal sections. (**B**) Representative photomicrographs in two different magnifications showing c-Fos expression in three sections of the IO (demarcated by blue dotted lines). (**C**) Representative photomicrographs showing c-Fos expression in two sections of the LRN (demarcated by red dotted lines). RDk: rostral nucleus of Darkschewitsch; fr: fasciculus retroflexus; MA3: accessory oculomotor nucleus; Dk: nucleus Darkschewitsch; D=dorsal; L=lateral.

**Figure 5—figure supplement 1.**
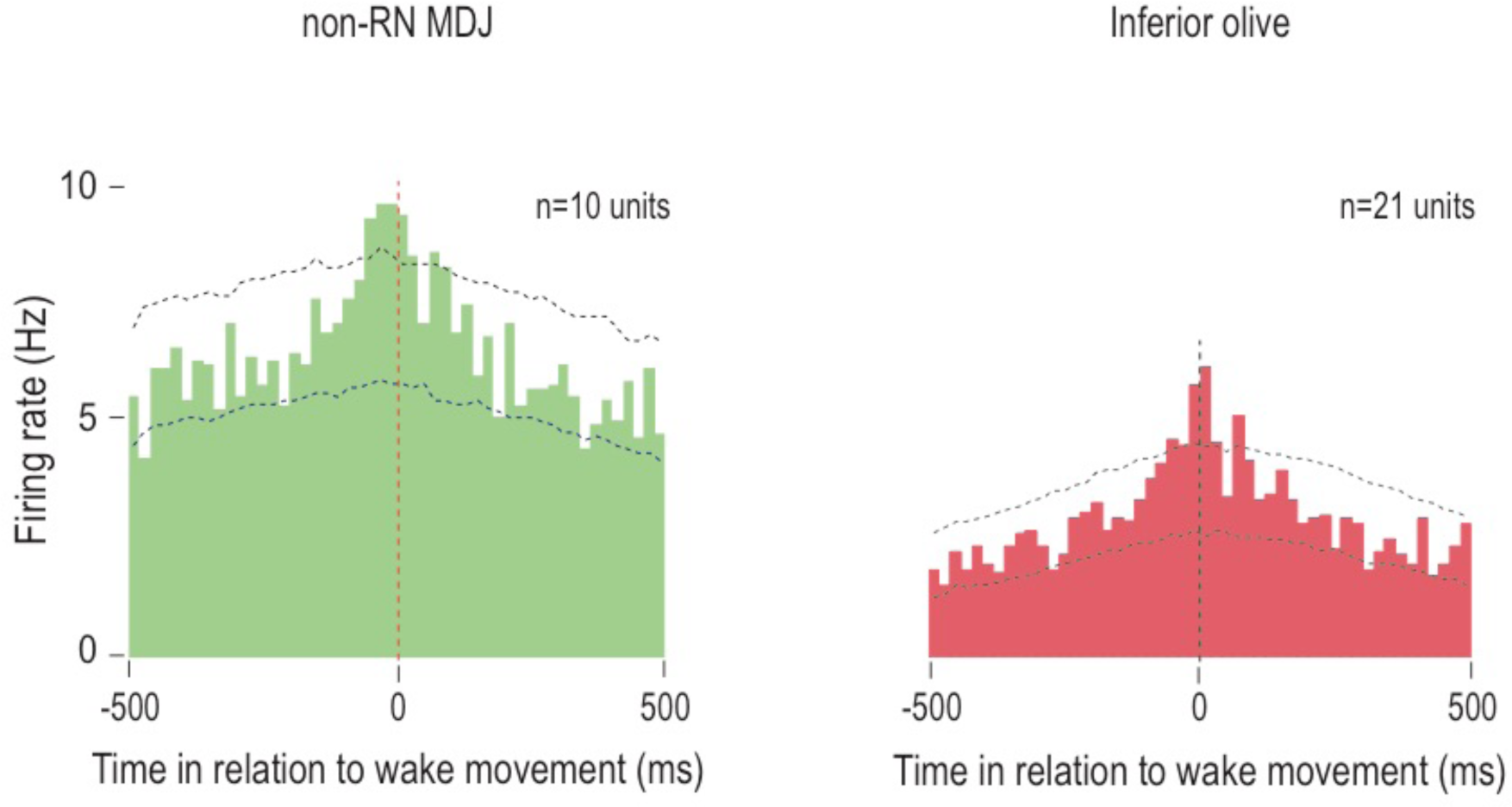
(**A**) Perievent histogram showing activity of twitch-preceding MDJ units (outside the RN) in relation to wake movements. Data pooled across 10 units and triggered on 586 wake movements. (**B**) Perievent histogram showing IO unit activity in relation to wake movements for those units that were significantly active in the ±10-ms time window around a twitch. Data pooled across 21 units and triggered on 747 wake movements.

**Figure 6—figure supplement 1.**
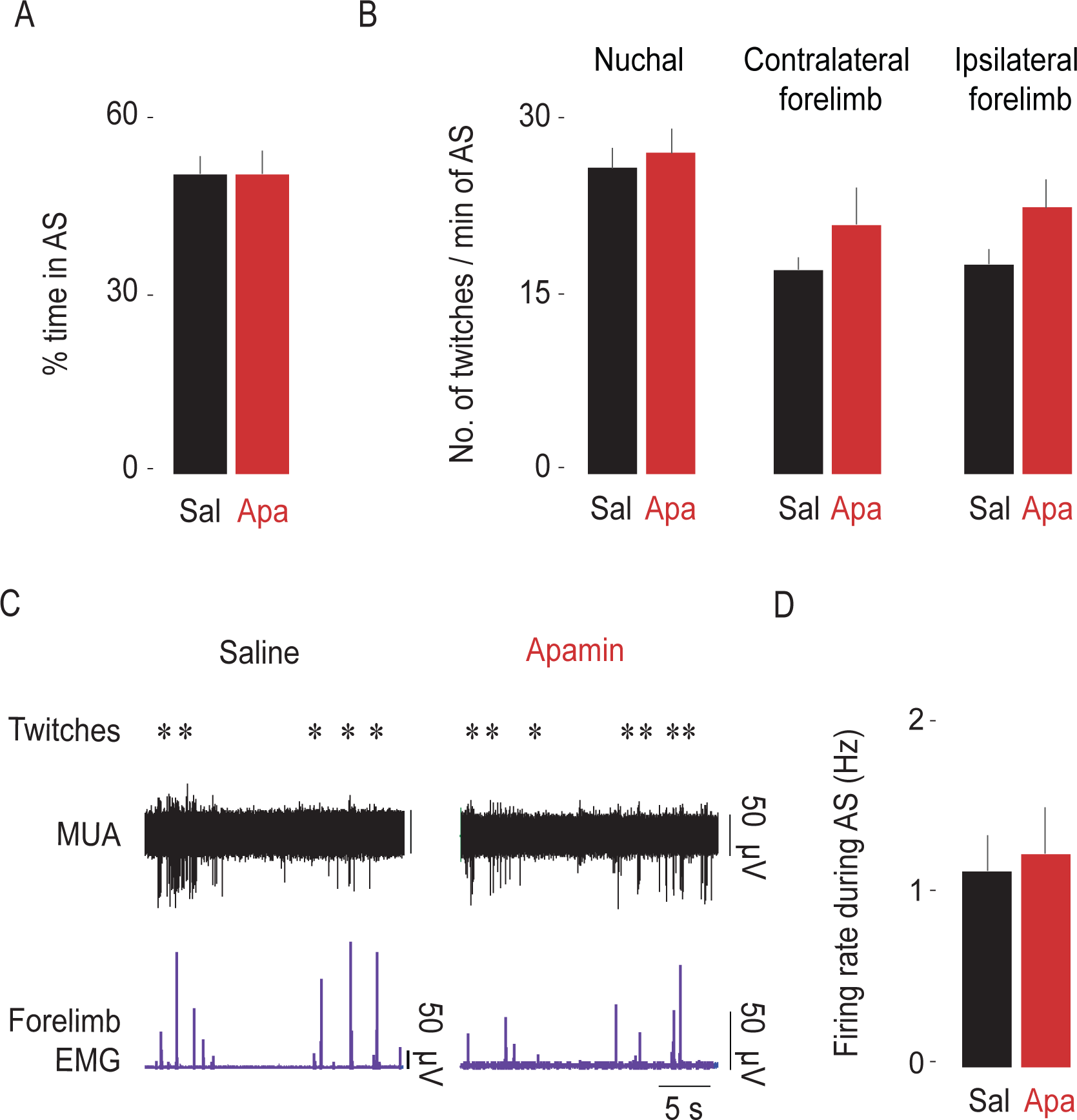
Apamin does not affect sleep-wake behavior. (**A**) Bar graphs showing the mean (+SEM) percentages of time spent in active sleep in saline (n=10 pups) and apamin groups (n=8 pups). (**B**) Bar graphs showing the mean (+SEM) rates of nuchal, contralateral and ipsilateral forelimb twitching in saline (n=10 pups) and apamin (n=8 pups) groups. (**C**) Representative recording of rectified forelimb EMG activity, and MUA in the IO during spontaneous sleep-wake cycling in saline (left) and apamin (right) groups. Asterisks denote twitches and gray horizontal bars denote wake movements as scored by the experimenter. (**D**) Bar graphs showing the mean (+SEM) firing rates of units during AS in saline (n=18 units) and apamin groups (n=22 units). Sal=saline; Apa=apamin.

## Acknowledgments

The authors thank Alex Tiriac and Jimmy Dooley for helpful comments. Research was supported by grants from the National Institute of Health (R37 HD-081168 and R01 HD-063071) to MSB. The authors declare no competing financial interests.

